# Rapid Assessment of Immune Effector Cell-mediated Cytotoxicity using mRNA Lipid Nanoparticles

**DOI:** 10.64898/2026.01.24.701542

**Authors:** Philip Mollica, Jian Li, Sang-Hoon Kim, Yingshi Chen, Aditya Bhartiv, Dillon O’Neil, Evan Weber, Mohamad Alameh, Leyuan Ma

## Abstract

Cellular immunotherapy has revolutionized cancer treatment by enabling more targeted and personalized disease management. As the field progresses, there is an increasing need for high-throughput in vitro assays to efficiently assess the cytotoxicity of therapeutic cells. Conventional cytotoxicity assays pose various limitations in the workflow and scalability. Here, we present an mRNA lipid nanoparticle (mRNA-LNP) approach to efficiently and robustly deliver reporter genes to target cells for assessing immune effector cell-mediated cytotoxicity. This approach enables the rapid, homogenous reporter expression without compromising the viability of target cells. The cytotoxicity results obtained using mRNA-LNP-transfected cells are highly consistent and comparable to those obtained using cell lines with stable reporter gene expression. Finally, we highlight the mRNA-LNP approach’s compatibility across a diverse range of tumor models, including primary tumor-derived models, enabling rapid and high-throughput assessment of the potency of various cytotoxic therapeutic cells.

## Introduction

The advances of cellular immunotherapy have revolutionized oncology, shifting the paradigm from broad-spectrum treatments toward personalized therapies^1,2^. The adoptive transfer of T cells engineered with chimeric antigen receptors (CARs) has demonstrated profound control of hematological malignancies such as relapsed and refractory B-cell acute lymphoblastic leukemia (B-ALL), lymphoma, and multiple myeloma^3,4^. The recent US-FDA approval of T cell receptor (TCR)-engineered T cells and tumor-infiltrating lymphocyte (TIL) therapies, coupled with numerous clinical trials investigating alternative immune effector cells such as natural killer (NK) cells, signals a promising and expanding future for cellular immunotherapies^5–7^.

An integral part of cellular immunotherapy development remains the assessment of therapeutic cells’ cytotoxic potency^8^. Such potency evaluations are needed throughout the lifecycle of a cellular therapy product, from early-stage candidate selection and mechanistic studies to late-stage product quality control^9^. Traditional cytotoxicity assays measure target cell cytolysis through the release of compounds such as Chromium-51^10^. Today, gold standard assays rely on target cells virally transduced to express reporter proteins such as GFP or firefly Luciferase(Fluc)^11^. However, many tumor cell lines, especially primary cells^12^, are difficult to transduce, requiring flow sorting, antibiotic selection, or single-cell cloning to generate homogeneous positive reporter lines. These processes are time-consuming and sometimes can reduce valuable tumor cell heterogeneity, thereby limiting the clinical relevance of the resulting data. Additionally, viral transduction poses the risk of insertional mutagenesis, which can alter target cell phenotype^13^. Transiently delivering reporter-encoding nucleic acid, such as DNA or RNA, using transfection or electroporation has been utilized for generating reporter cells. However, these approaches are often costly, require optimization for each cell line, and often severely reduce cellular viability^14^. Recognizing these limitations, we sought to develop a faster, safer, and more robust approach for generating reporter cell lines that is compatible with the broad spectrum of tumor models and immune effector cell types.

Here, we demonstrate that LNP delivery of mRNA encoding a reporter gene, GFP or FLuc, to target cells enables the rapid assessment of immune effector cell cytotoxicity (**Figure.1A**). The entire process, including target cell transfection, co-culture and cytotoxicity readout, can be completed as soon as eight hours. The mRNA-LNP-based cytotoxicity assay yielded results that are highly consistent with those obtained using stable reporter cell lines, offering a highly reliable, versatile and cost-effective approach for rapidly assaying immune cell cytotoxicity.

## Materials and Methods

### Tumor Cell Lines and Cell Culture

NALM6, RD, and MDA-MB-231, and B16F10 cells were purchased from ATCC. The primary tumor-derived human cell lines CY1006, JL27, KL1118, and BT1018 were kindly provided by the Cancer Cell Line Factory of the Broad Institute at MIT. The primary tumor cell line SJSA-1 was kindly provided by the Mitochondrial Medicine Program of the University of Pennsylvania. The cell lines NK92 and A375 were kindly provided by Dr. Xiaowei Xu from the University of Pennsylvania. NALM6 cells were maintained in RPMI (Cytiva) media supplemented with 10% FBS (Sigma-Aldrich) and 1% penicillin-streptomycin (Gibco). NK92 cells were cultured in MEM-α media supplemented with 10% FBS, 1% penicillin-streptomycin, hIL2 (PeproTech) (100 IU/ml), 1% 2-mercaptoethanol (Gibco), and 10% horse serum (Gibco). All other tumor cell lines were cultured in DMEM (Cytiva) supplemented with 10% FBS and 1% penicillin-streptomycin. All cells were cultured at 37 °C and 5 % CO_2_.

### Human CAR T cell Preparation and Culture

Primary human T cells were obtained from the Human Immunology Core at the University of Pennsylvania and transduced as before^15^. Briefly, T cells were activated using CD3/CD28 Dynabeads (Gibco) for 24 hours and kept in culture throughout transduction. Transduction was achieved by co-culturing T cells with lentiviral supernatant and polybrene (Millipore Sigma) (1:2000), and centrifuging them at 2000 G for 90 mins at 35 °C. These cells were then expanded until ready to use, usually till day 10. Cells were maintained in RPMI supplemented with 10% FBS and hIL2 (Peprotech) (100 IU/ml), and cultured at 37 °C and 5 % CO_2._ CAR expression was confirmed using flow cytometry.

### Murine CAR T cell Preparation and Culture

Primary murine T cells were isolated from the spleen of a C57BL/6 mouse using the EasySep CD3 Isolation kit (Stem Cell Technologies) and transduced as previously described^15,16^. Briefly, cells were immediately cultured on a non-TC treated 6 well plate (Falcon) pre-coated with the murine anti-CD3 (2C11, 0.5μg/ml) and anti-CD28 (37.51, 5μg/ml) antibody (Bio X Cell) for 48 hours. Following activation, transduction was performed by mixing cells with retroviral supernatant and polybrene (1:2000), and centrifuging them at 2000 G for 90 mins at 35 °C. These cells were then expanded until ready to use, usually on day 5 post-activation. Cells were maintained in RPMI supplemented with 10% FBS, 1X Non-Essential Amino Acids (Gibco), 1X 2-mercaptoethanol (Gibco), 1X Sodium Pyruvate (Gibco), 1% Insulin-Transferrin-Selenium (Gibco), and mIL2 (10 ng/ml), and cultured at 37 °C and 5 % CO_2._ CAR expression (myc tag) was confirmed using flow cytometry.

### Production of mRNA

The mRNA encoding codon-optimized sequences of the firefly luciferase (FLuc) or the eGFP (eGFP) were purchased from RNA technologies (Montreal, Canada). The preclinical grade mRNA was produced through co-transcriptional capping using the CleanCap analogue, in the presence of 1N methylpseudouridine, ribonucleotide phosphates (rNTPs), and a T7 polymerase. The reaction was performed using an in-house optimized buffer proprietary to RNA technologies. The DNA template was digested using DNAse I, and the mRNA product underwent purification and buffer exchange. The purity of the final product was assessed with spectrophotometry, capillary gel electrophoresis, and dot blot (dsRNA). Endotoxin content was measured using a chromogenic Limulus amebocyte lysate (LAL) assay; all assays were negative.

### LNP preparation

LNP were formulated by microfluidic mixing as previously described^17^. Briefly, SM102, DSPC, cholesterol, and DMG-PEG2000 were mixed in the ethanol phase at molar ratios of 50:10:38.5:1.5 and rapidly mixed with the aqueous phase containing either FLuc or eGFP mRNA. The formed LNPs were concentrated using 10 kDa Amicon Ultra 15 mL centrifugal filter units (Millipore Sigma), and frozen at −80 °C until further use. The size and PDI were determined via dynamic light scattering (DLS) using the Zetasizer Ultra (Malvern Panalytical). mRNA encapsulation efficiency and quantification were determined using the Quant-it Ribogreen reagent (Thermo Fisher, R11491).

### Flow Cytometry

All flow cytometry experiments were performed using the BD LSRII cytometer and analyzed using FlowJo v10 software. Human CAR expression was assessed using AF647-conjugated anti-Myc tag (Cell Signaling Technologies) or APC-conjugated anti-flag tag antibodies (Biolegend). Murine CAR expression was assessed using an AF647-conjugated anti-myc tag antibody(Cell Signaling Technologies). GFP expression was confirmed by reading the fluorescence following excitation at 480 nm and passed through the 530/30 filter. Positive gating was determined relative to the autofluorescence of each respective tumor cell line. 7-aminoactinomycin D (Biolegend) was used to label dead cells across all flow cytometry experiments. EGFRvIII expression by KL1118 was confirmed using an anti-EGFRvIII scFv-Fc fusion protein^15^ followed by an PE conjugated anti-Fc (Biolegend). OVA peptide-loaded MHCI expression by B16F10 cells following SIINFEKL peptide pulsing was confirmed using PE-conjugated anti-H2Kb (BD Biosciences).

### Incucyte Cytotoxicity Assay

Kinetics of immune cell-mediated target cytolysis were quantified using the image-based Sartorius Incucyte system. 2.5×10^4^ target cells expressing GFP were plated with effector cells at 5/2/1/0.5:1 Effector:Target (E:T) ratios. Solid tumor models were allowed to adhere prior to plating effector cells. Total culture volume was 200 μl in a 96 well flat bottom plate. Plates were imaged at 10X zoom with 5 images per well. Photos were taken every hour of co-culture and analyzed for 24 hours until the experimental endpoint. Tumor cells were tracked in terms of Green Total Integrated Intensity (TII). For human and murine CAR T cell models, data was normalized to target wells cultured with non-transduced T cells. For the NK cell assay, data was compared back to target cells cultured alone. Percent cytolysis was quantified using the equation:

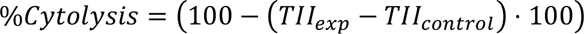

where TII_exp_ = experimental group and TII_control_ = control group.

### Luciferase Cytotoxicity Assay

NALM6 cells were transfected with 1 µg of luciferase mRNA–LNP per million cells and incubated for 4-24 hours. After transfection, cells were washed once with medium and plated in a white plate with varying numbers of either untransduced T cells or CD19 CAR-T cells at defined Effector:Target (E:T) ratios. At the end time point, Luciferin (Revvity) was added at a final concentration of 150 μg/ml. The plate was immediately placed into a Tecan Spark Microplate Reader for analysis of total luminescence. Data was normalized to target cells cultured with non-transduced T cells. Percent cytolysis was quantified using the equation provided above.

### Statistics

To directly compare the cytotoxicity data generated by target cells produced by either mRNA-LNP transfection or lentiviral (LV) transduction, we first plotted the killing kinetics for each group at each respective effector-to-target ratio. We then pooled the data and generated a line of best fit based on an exponential plateau model. The fitness of this line for the overall data set was evaluated in terms of the coefficient of determination (r^2^). The coefficient of determination assesses the variation in the dependent variable (% Cytotoxicity) when the independent variable is changed. In our experiments, the independent variable is the method of generating the reporter cell line used in the assay: mRNA-LNP vs LV. The high r^2^ we observed demonstrates that there is minimal variability in the data, regardless of the approach used to generate the fluorescent reporter cell line. All statistical analysis was performed using GraphPad Prism v10.

## Results

### mRNA-LNPs enable efficient and non-toxic target cell labeling and produce comparable results to virally transduced cells in cytotoxicity assays

To investigate the ability of mRNA-LNPs to rapidly generate reporter cell lines for use in cytotoxicity assays, we first standardized this approach using GFP mRNA-LNPs and the well-established NALM6 B-cell acute lymphoblastic leukemia model^18^. To determine the kinetics of GFP expression following mRNA-LNP transfection, NALM6 cells were incubated with GFP mRNA-LNPs at various concentrations. Flow cytometry analysis showed that GFP mRNA-LNPs rapidly transfected NALM6 cells, with nearly 100% of cells expressing GFP in the first 4 hours post-transfection and peak GFP expression at 24 hour (**Figure. 1B**). This expression pattern is consistent across all mRNA-LNP concentrations tested and homogenous GFP expression was noted to remain stable for up to 72 hours (**Figure. 1B-C**). Notably, LNP transfection was non-toxic, and the viability of transfected cells was maintained above 85% across all doses and timepoints (**Figure.1D**).

**Figure 1:**
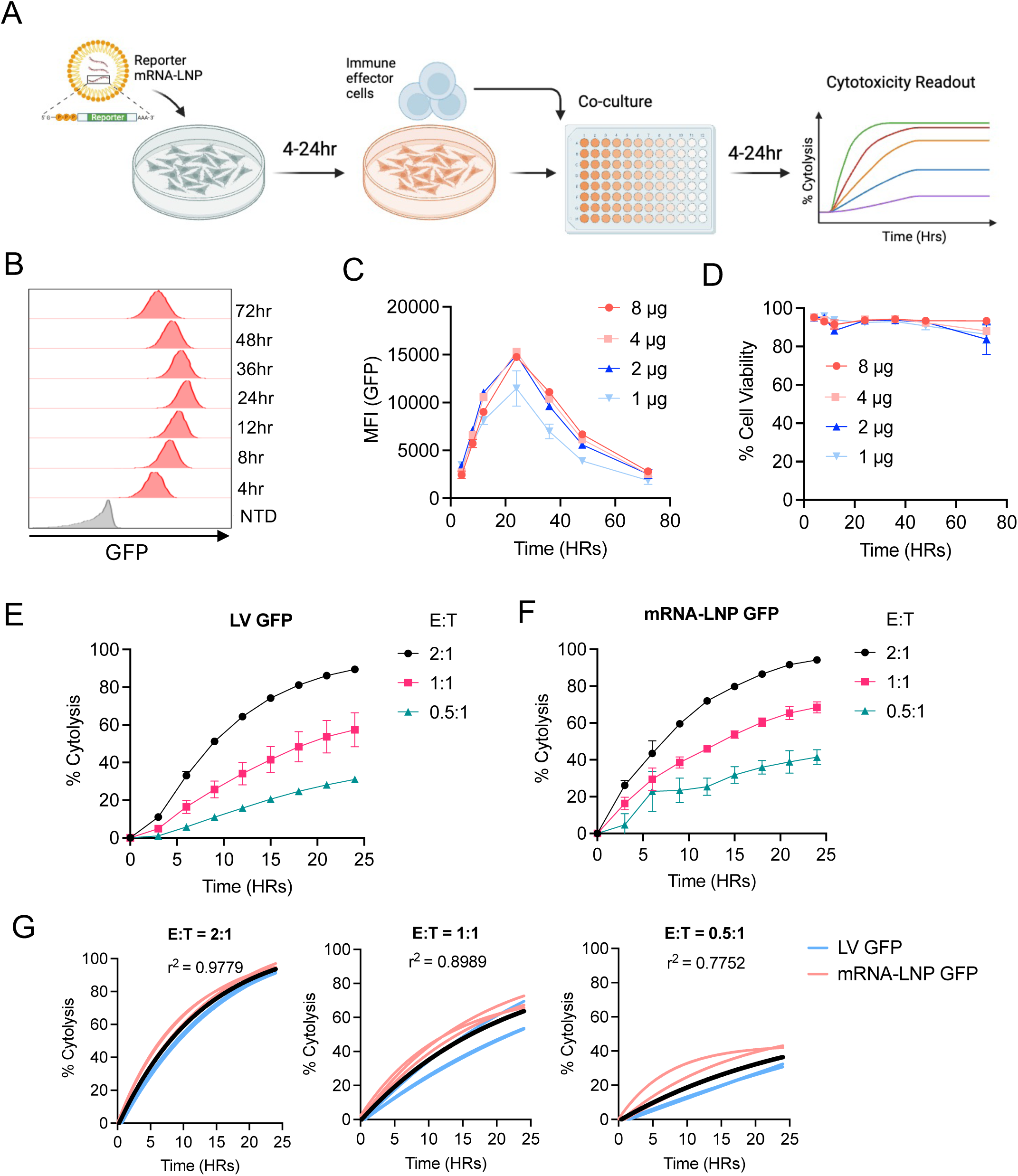
mRNA-LNP reporter assay enables robust assessment of CAR-T cytotoxicity. (A) Schematic timeline illustrating the mRNA-LNP reporter delivery and downstream cytotoxicity assay. (B) Representative flow cytometry histograms highlight the kinetics of GFP expression in NALM6 cells following GFP mRNA-LNP transfection at [1 µg / 10^6^ cells]. Wildtype NALM6 cells were used as a control. (C) GFP expression by NALM6 cells post-transfection by different doses of GFP mRNA-LNP over time. Shown was the geometric mean fluorescent intensity (gMFI). (D) Trending viability of NALM6 cells post-transfection of mRNA-LNP from (C). (E-F) CD19-CAR T cytotoxicity data generated using Incucyte following 24-hour co-culture with GFP^+^ NALM6 cells generated by LV transduction (E) or mRNA-LNP transfection (F). Target cells cultured with non-transduced donor-matched T cells served as a negative control. (G) Coefficient of determination (r^2^) for the cytotoxicity data using LV or LNP-generated GFP^+^ NALM6 cells from each indicated E:T ratio. For all cytotoxicity data, lines of best fit were generated using nonlinear fit within GraphPad Prism. Error bars show mean ± SD (n=3). Graph in (A) was generated using biorender.com

To compare the utility of GFP mRNA-LNP approach against the standard viral transduction approach, NALM6 reporter cells generated using either approach were co-cultured with CD19-CAR T cells or non-transduced human T cells at various effector-to-target (E:T) ratios. Kinetics of tumor cell killing were monitored by Incucyte for 24 hours. Across all settings, we observed comparable killing kinetics and a dose-dependent responsiveness between LV (**Figure. 1E**) and mRNA-LNP generated target cells (**Figure. 1F**), evidenced by the high and significantly uniform coefficient of determination (r^2^) across all groups (**Figure.1G**).

### The mRNA-LNPs approach enables rapid assessment of cytotoxicity in the same day

To demonstrate the versatility of the mRNA-LNP cytotoxicity approach, we used FLuc as an alternative reporter. Upon transfection with FLuc mRNA-LNPs, NALM6 cells exhibited a rapid increase in luminescent intensity with the signal peaks at 24 hours post-transfection and slowly declined till 72 hours (**Figure.2A**), a trend similar to that from GFP mRNA-LNP (**Figure. 1B**). We then co-cultured the mRNA-LNP or LV-derived FLuc-expressing NALM6 cells with CD19-CAR T cells at various E:T ratios and incubated for 24 hours. The cytotoxicity data confirmed a dose-dependent cytotoxicity and were highly comparable across LV and mRNA-LNP groups with a coefficient of determination (r^2^) of 0.8864 (**Figure. 2B**).

**Figure 2:**
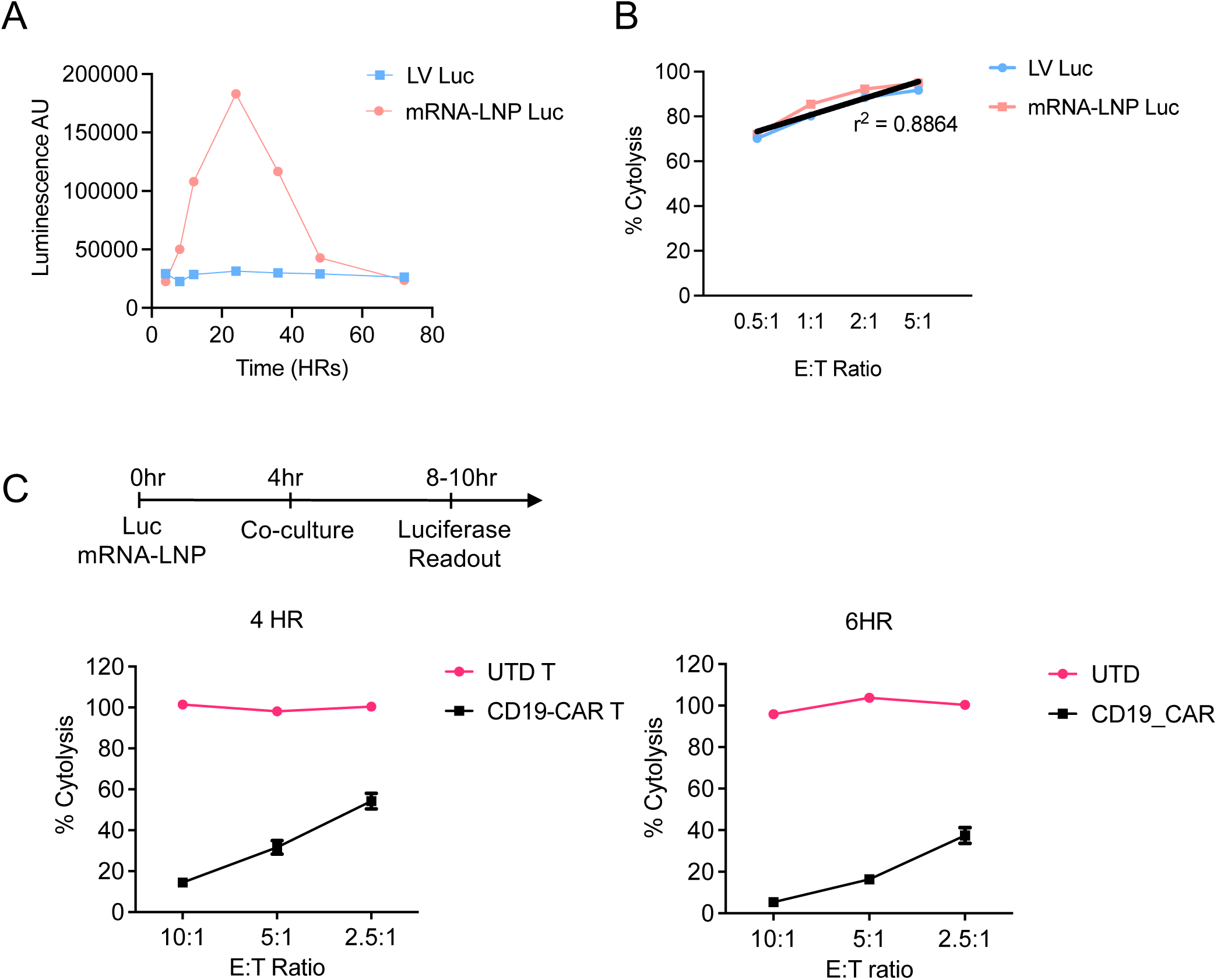
Rapid assessment of CAR-T cytotoxicity using the FLuc-mRNA-LNP reporter assay. (A) Kinetics of luciferase expression in NALM6 cells following FLuc mRNA-LNP transfection. NALM6 cells stably transduced with FLuc-encoding LV was included as control. (B) Cytotoxicity assessment using LV-Luc or mRNA-LNP Luc generated reporter NALM6 cells co-culturing with CD19-CAR T cells. Target and effectors were co-cultured for 24 hours prior to measuring luminescence. Gross cytotoxicity data were compared across the LV and LNP-generated FLuc^+^ NALM6 cells, coefficient of determination (r^2^) was generated within GraphPad Prism using simple linear regression. (C) An accelerated assessment of cytotoxicity using FLuc mRNA-LNP reporter delivery. Shown is the timeline. NALM6 cells were transfected with FLuc mRNA–LNP for 4 hours, then cytotoxicity data were collected after 4hour or 6 hour of co-culture with either untransduced T cells (UTD) or CD19 CAR-T cells at the indicated E:T ratios. ***, p<0.001 by student’s t tests.

Given the sensitivity of the luciferase assay, we next sought to test if the FLuc-mRNA-LNP approach allows the cytotoxicity assay to be completed in the same day. NALM6 cells were transfected with FLuc-mRNA-LNP for 4 hours, then harvested and co-cultured with CD19-CAR T cells or untransduced T cells for 4 or 6 hours at various E:T ratios (10:1, 5:1 or 2.5:1). Specific cytotoxicity could be robustly detected across all conditions, with the 10:1 E:T ratio yielding >80% specific target cell lysis as early as 4 hours after co-culture (**Figure. 2C**).

Together, these data highlight the robustness, sensitivity, and efficiency of using FLuc-mRNA-LNPs to evaluate CAR T-cell cytotoxicity, which can be completed in as few as 8 hours.

### mRNA-LNPs approach for assessing cytotoxicity against established and primary solid tumors

We next sought to assess the mRNA-LNP-based cytotoxicity approach in solid tumor models. To this end, B16F10 murine melanoma cells were transfected with GFP mRNA-LNP at various doses. Flow cytometry analysis confirmed stable GFP expression throughout the 48-hour observation period at an expression level comparable to that achieved with the LV transduction approach (**Figure. 3A**). Greater than 90% cell viability was maintained across all time points and LNP doses (**Figure. 3B**). Next, mRNA-LNP and LV-based reporter B16F10 cells (TYRP1^+^) were co-cultured with TYRP1-targeting mouse CAR T cells^15^ at various E:T ratios for 24 hours and cytotoxicity was monitored using Incucyte. Both reporter cells exhibited nearly identical cytotoxicity kinetics (**Figure.3C-D**) and dose-dependent responsiveness with r^2^ >0.9 across all conditions (**Figure. 3E**).

**Figure 3:**
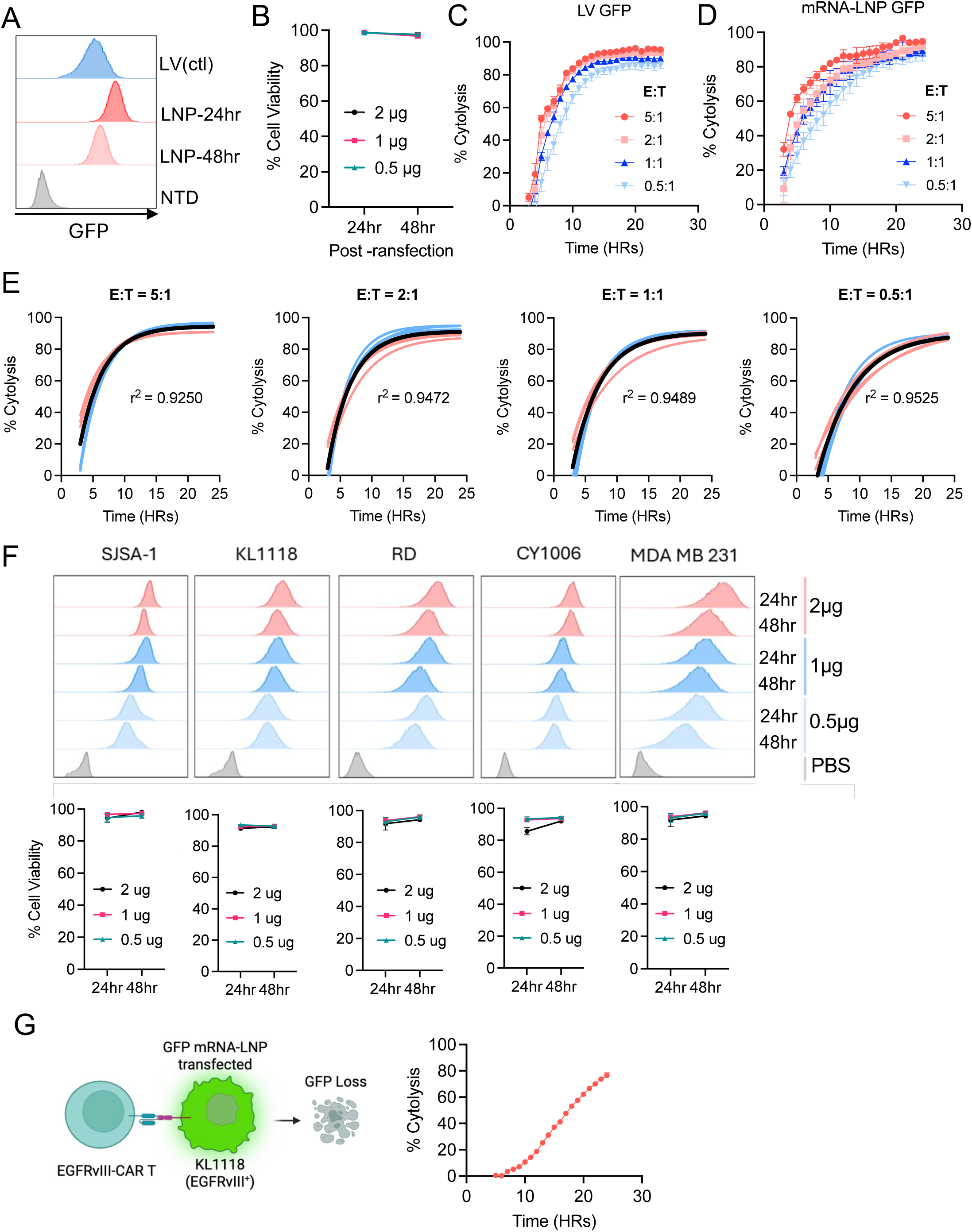
The mRNA-LNP reporter assay is compatible with solid tumor and primary tumor models for CAR-T cytotoxity assessment. (A) Representative flow cytometry histograms showing GFP expression of both the LV and mRNA-LNP generated B16F10 target cells. (B) Viability data showing LNP transfection was well tolerated by B16F10 cells. (C-D) Anti-TYRP1-CAR T cytotoxicity data generated using Incucyte following 24-hour co-culture with GFP^+^ B16F10 cells generated by LV transduction (C) or mRNA-LNP transfection (D). Target cells cultured with non-transduced donor-matched T cells served as a negative control. (E) Coefficient of determination (r^2^) for the cytotoxicity data using LV or LNP-generated GFP^+^ B16F10 cells from each indicated E:T ratio. (F) Representative histograms and viability data following transfection of a variety of solid tumor cell lines with GFP mRNA-LNPs. (G) Representative schematics and Incucyte cytotoxicity data generated using EGFRvIII-CAR T cells co-cultured with GFP^+^ KL1118 cells generated by GFP mRNA-LNP transfection. Error bars show mean ± SD (n=3). Graph in (G) was generated using biorender.com

We next sought to test if the mRNA-LNP cytotoxicity approach is compatible with patient-derived primary tumor models, as they reliably represent tumor biology and are frequently employed to test the functionality of immunotherapies^19,20^. Notably, these cell lines are often difficult to transduce using viral vectors^21^. These patient-derived tumor models were selected to encompass the broad spectrum of commonly studied tumor types: osteosarcoma (SJSA-1), melanoma (CY1006), glioblastoma (KL1118). We also included two additional established cell lines, rhabdomyosarcoma (RD) and triple-negative breast cancer (MDA-MB-231), to ensure the generalizability. All cell lines were transfected with GFP mRNA-LNPs and exhibited homogeneously high and stable GFP expression over 48 hours, with cell viability exceeding 80% (**Figure 3F**). We next used GFP mRNA-LNP-transfected EGFRvIII^+^ KL1118 cells as the target cells to assess the cytotoxicity of human EGFRvIII-targeting CAR T cells^15,16^. We found that EGFRvIII-CAR T cells could progressively kill EGFRvIII^+^ KL1118 cells up to 80% within 24 hours at a 5:1 E:T ratio (**Figure. 3G**).

Overall, these data demonstrate the capacity of GFP-LNPs to enable the high-throughput generation of reporter cells for rapid assessment of T-cell-mediated cytotoxicity in clinically relevant tumor models.

### The mRNA-LNP-based cytotoxicity assay is broadly applicable for assessing immune effector cell-mediated target cell killing

To expand the scope of our study, we validated the use of GFP mRNA-LNPs to measure NK cell-mediated cytotoxicity using the well-established NK92 cell line^22^. The human melanoma cell line A375 is a known target of NK92 and was used for this experiment^23^. Consistent as before, transfection of A375 cells with GFP mRNA-LNP led to robust GFP expression across a 48-hour observation period (**Figure. 4A**). The mRNA-LNP-transfected or LV-transduced A375 cells were then co-cultured with NK92 cells at various E:T ratios and analyzed for cytotoxicity in Incycyte for 24-hour (**Figure. 4B-C**). The results demonstrated a strong consistency in terms of dose responsiveness, killing kinetics, and overall cytotoxicity between the mRNA-LNP and LV groups across all E:T ratios with an r^2^ > 0.86 to monitor NK cell-mediated cytotoxicity(**Figure 4D**).

**Figure 4:**
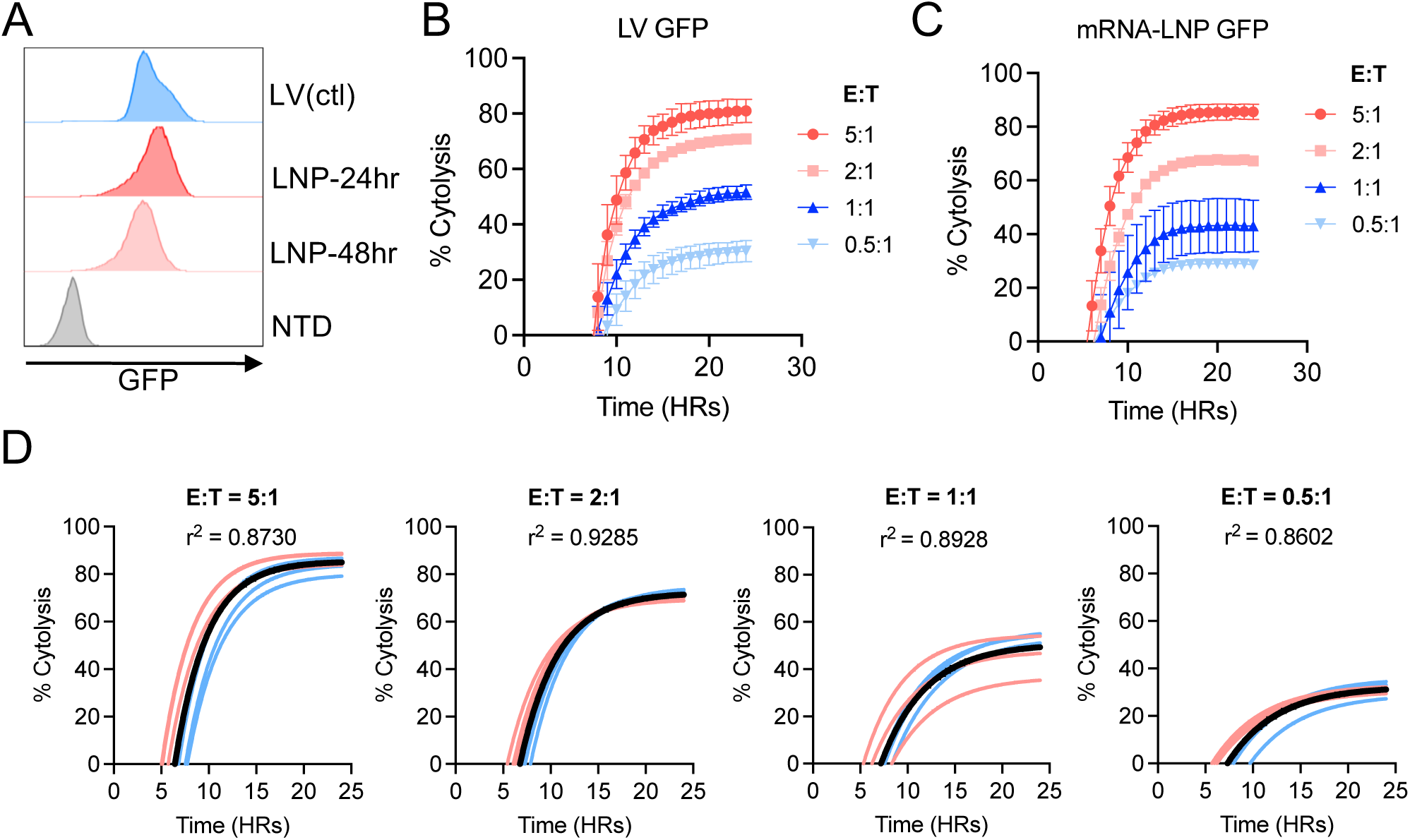
The mRNA-LNP reporter assay is compatible with NK cell-mediated cytotoxicity. (A) Representative flow cytometry histograms of the GFP expression of both the LV and LNP generated A375 human melanoma cells. (B-C) NK92 cytotoxicity data generated following 24 hour co-culture with GFP^+^ A375 cells produced by LV transduction (B) or mRNA-LNP transfection (C). Target cells cultured without NK92 cells served as a negative control. (D) Coefficient of determination (r^2^) for the cytotoxicity data using LV or LNP-generated GFP^+^ A375 cells from each indicated E:T ratio. For all cytotoxicity data, lines of best fit were generated using nonlinear fit within GraphPad Prism. Error bars show mean ± SD (n=3).

## Discussion

A major bottleneck in the development lifecycle of cell-based immunotherapies is the screening of lead candidates to assess their relative therapeutic potency. Complicating these efforts is the use of standardized biological assays, which can be time-consuming and challenging to scale.In this study, we address these concerns by introducing a streamlined protocol leveraging mRNA-LNP to transiently generate reporter cell lines, enabling rapid and high-throughput evaluation of therapeutic immune effector cells’ cytotoxicity. Our method ensures homogenous and nontoxic reporter protein expression without the need for time-consuming optimization. Notably, this entire protocol can be completed for a desired target cell line in as few as eight hours, including both reporter gene delivery using mRNA-LNPs and cytolysis by immune effector cells. Additional strengths lie in the flexibility of reporter-gene selection and in highly consistent efficacy across a broad range of established and patient-derived tumor models. Importantly, data generated using the mRNA-LNP approach were significantly similar to those from reporter systems generated by the gold-standard LV transduction across multiple immune effector cell types, including both CAR T and NK cells. Together, these properties make the mRNA-LNP-based cytotoxicity assay a generalizable and robust alternative for determining the potency of immune effector cells and aiding the development of various cellular immunotherapies.

## Acknowledgements

We thank Dr. Xiaowei Xu of the University of Pennsylvania for sharing NK92 and A375 cells. We thank the Mitochondrial Medicine Program of the University of Pennsylvania for sharing SJSA-1 cells. We thank the Cancer Cell Line Factory of The Broad Institute at MIT for the patient derived glioblastoma cell line KL1118. Funding: The work was supported by the Cell and Gene Therapy Collaborative and the Junior Faculty Pilot Program at CHOP, NIH NEW Innovators Award (DP2 AI164319-03), ITMAT at Penn and NCATS (UL1TR001878).

## Data and Materials Availability

All data are available on request.

## Author Contributions

P.M., J.L. S.K., and L.M. designed the experimental strategy. P.M. and L.M. wrote the manuscript. P.M., J.L. S.K., Y.C., and A.B. performed the experiments. D.N. and M.G.A. provided all mRNA-LNPs. P.M. and L.M. analyzed and interpreted the data. P.M. J.L. and L.M. prepared the figures. All authors reviewed and commented the manuscript.

